# DNMT3A harboring leukemia-associated mutations directs sensitivity to DNA damage at replication forks

**DOI:** 10.1101/2021.05.28.445639

**Authors:** Kartika Venugopal, Pawel Nowialis, Yang Feng, Daniil E. Shabashvili, Cassandra M. Berntsen, Kathryn I. Krajcik, Christina Taragjini, Zachary Zaroogian, Heidi L. Casellas Román, Luisa M. Posada, Chamara Gunaratne, Jianping Li, Daphné Dupéré-Richer, Richard L. Bennett, Santhi Pondugula, Alberto Riva, Christopher R. Cogle, Rene Opavsky, Brian K. Law, Stefan Kubicek, Philipp B. Staber, Jonathan D. Licht, Jonathan E. Bird, Olga A. Guryanova

## Abstract

Mutations in the DNA methyltransferase 3A (*DNMT3A*) gene are recurrent in *de novo* acute myeloid leukemia (AML) and are associated with resistance to standard chemotherapy, disease relapse, and poor prognosis, especially in advanced-age patients. Previous gene expression studies in cells with *DNMT3A* mutations identified deregulation of cell cycle-related signatures implicated in DNA damage response and replication fork integrity, suggesting sensitivity to replication stress. Here we tested whether pharmacologically-induced replication fork stalling creates a therapeutic vulnerability in cells with *DNMT3A*(R882) mutations. We observed increased sensitivity to nucleoside analogs such as cytarabine in multiple cellular systems expressing mutant *DNMT3A*, ectopically or endogenously, *in vitro* and *in vivo*. Analysis of DNA damage signaling in response to cytarabine revealed persistent intra-S phase checkpoint activation, accompanied by accumulation of DNA damage in the *DNMT3A*(R882) overexpressing cells, which was only partially resolved after drug removal and carried through mitosis, resulting in micronucleation. Pulse-chase double-labeling experiments with EdU and BrdU after cytarabine wash-out demonstrated that cells with *DNMT3A(mut)* were able to restart replication but showed a higher rate of fork collapse. Gene expression profiling by RNA-seq identified deregulation of pathways associated with cell cycle progression and p53 activation, as well as metabolism and chromatin. Together, our studies show that cells with *DNMT3A* mutations have a defect in recovery from replication fork arrest and subsequent accumulation of unresolved DNA damage, which may have therapeutic tractability. These results demonstrate that, in addition to its role in epigenetic control, DNMT3A contributes to preserving genome integrity during DNA replication.

## INTRODUCTION

Acute myeloid leukemia (AML), a genetically heterogeneous malignant clonal disorder of the hematopoietic system, is the most common leukemia in adults^1,2^. Its incidence sharply increases with age, with median age at diagnosis 67-68 years. Due to intensive basic and clinical research, the long-term survival has significantly increased in the last 50 years and is currently approaching 30%-40%. Development of targeted therapies has led to remarkable successes in some genetic subtypes of AML leading to improved survival and quality of life. However, for most AML subtypes targeted approaches are still unavailable, leaving cytotoxic chemotherapy the next best option^3,4^. To date, therapeutic targeting for AML with *DNMT3A* mutations has not shown much promise.

Recurrent mutations in the DNA methyltransferase 3A (*DNMT3A*) gene are found in 20-35% of *de novo* AMLs and are associated with poor prognosis^5-12^, in part attributable to relative resistance to anthracyclines^12-14^. While chemotherapy dose intensification demonstrated some survival benefit^12,14^, outcomes remain unsatisfactory in most patients with *DNMT3A* alterations due to advanced age, comorbidities, and inability to tolerate treatment^1,7,15^. Cytarabine (Ara-C) remains a foundation of AML treatment^16^, with low-dose Ara-C being a strategy of choice in advanced-age patients^3,17-19^. Clinical trials of low-intensity regimens combining cytarabine and cladribine, nucleoside analog chain terminators that stall DNA replication, with and without venetoclax, are safe and effective^3,20^, and in a recent study tended to benefit patients with *DNMT3A* mutations^21^. The most common mutation type, representing up to 70% of all *DNMT3A* alterations in AML, is single amino-acid substitution at arginine 882 (R882, most commonly R882H or R882C) in the catalytic domain^9,11,12,22,23^. Hence, we and others performed DNA methylation analyses^13,24-27^ and gene expression profiling^13,28^ in primary AML samples and in animal models carrying *DNMT3A* mutations. These studies uncovered gene signatures of altered cell cycle control such as negative enrichment of the CHK1-regulated G2/M checkpoint^29,30^, which may indicate persistent replication stress, in cells expressing mutant *DNMT3A*^13,28^. Importantly, DNMT3A protein has been detected in direct association with stalled replication forks in cells treated with hydroxyurea^31^. Yet, its role in the context of replication fork stalling such as after cytarabine-based chemotherapy remains unknown.

Once cytarabine is converted into its active metabolite ara-CTP, it is incorporated into nascent DNA during replication where it is a poor substrate for chain extension^32^. This results in chain termination, replication fork stalling, and DNA damage response (DDR) often leading to p53-dependent apoptosis^33-36^. Single-stranded DNA breaks (SSBs) resulting from stalled replication forks actuate ATR-dependent CHK1 phosphorylation necessary to preserve replication fork integrity^37^. If unresolved, SSBs are converted to double-stranded DNA breaks (DSBs) and activate ATM-dependent phosphorylation cascade of CHK2, H2A.X, and p53, which in turn mediate DNA DSB repair, and cell cycle arrest and/or apoptosis^38-40^.

This study investigated the role of mutant DNMT3A (DNMT3A^mut^) in recovery of stalled replication forks and therapeutic responses to replication stress-inducing drugs. We show in multiple systems that cells expressing *DNMT3A*^*mut*^ are more sensitive to cytarabine-induced replication. This was accompanied by persistent intra-S checkpoint activation and accumulation of unrepaired DNA damage, which was carried through mitosis, leading to mitotic defects^41^. Mechanistic studies of replication fork dynamics in cytarabine-treated cells demonstrated a higher rate of replication fork collapse in cells with DNMT3A^mut^. These results provide mechanistic insights into the regulation of replication fork recovery after DNA damage, and suggest additional strategies to augment therapeutic responses to cytarabine and similar drugs used to treat AML.

## MATERIALS AND METHODS

### Cell lines

Human leukemia cell lines K-562, KU-812, SET-2, and KO-52 were grown in RPMI supplemented with 10% FBS and penicillin/streptomycin. U2OS cells were maintained in DMEM supplemented with 4.5 g/l glucose, nonessential amino acids, 10% FBS, and penicillin/streptomycin. Cells were lentivirally transduced to express wild-type or R882C-mutant DNMT3A or with empty vector control (pMIGR1 expressing GFP as a selectable marker, or pPICH expressing dsRED). For robustness, all experiments were performed with both sets of isogenic cell lines, one with pMIGR1 and one with pPICH, as independent biological replicates.

### Cell viability and drug dose response assays

Cells were plated in 96-well plates at 1×10^4^/well (suspension) and 3×10^3^/well (adherent) and exposed to cytarabine (Ara-C), fludarabine, or cladribine (Cayman Chemicals), or hydroxyurea (HU) (Sigma) in triplicate. Relative cell numbers were quantified at the indicated time points by CellTiter Glo (Promega) for suspension cultures or by alamarBlue (Thermo Fisher) for adherent cells. Plates were read with a BMG LABTECH FluoroQuant Optima plate reader. Relative cell viability data normalized to vehicle controls set at 100% were used to calculate IC_50_ values by fitting a four-parameter (variable slope) dose-response curve. GraphPad Prism version 7 software was used.

### Mice

All animal studies were approved by the University of Florida Institutional Animal Care and Use Committee. A knock-in mouse line that conditionally expresses *Dnmt3a*^*R878H*^ allele (equivalent to *DNMT3A*^*R882H*^ in humans) from the endogenous locus (Jackson Laboratory stock No. 031514) was described^13^. For inducible expression of the mutant form of *Dnmt3a* and generation of mouse leukemias with *Flt3*^*ITD*^ and *Npm1*^*c*^ alleles see Supplementary methods. For *in vivo* cytarabine treatment studies, pre-conditioned congenic CD45.1 recipients were transplanted with CD45.2 donor bone marrow from fully excised *Dnmt3a*^*+/R878H*^*:Mx1-Cre*^*+*^ (*Dnmt3a*^*mut*^) and littermate control *Dnmt3a*^*+/+*^*:Mx1-Cre*^*+*^ *(Dnmt3a*^*wt*^) mice. Once engrafted, animals were treated with 30mg/kg cytarabine/day for 5 days IP and vehicle (PBS) (*n*=5/group). Bone marrow was analyzed after 48h. See also Supplementary methods.

### Colony forming unit (CFU) assays

Freshly isolated whole bone marrow cells harvested from the *Dnmt3a* wildtype and mutant mice (described above) were plated in MethoCult M3434 (StemCell), 1×10^4^ cells/well in triplicate. Vehicle (PBS) or cytarabine were added directly to the methylcellulose media at indicated concentrations. Colonies were scored 10-14 days after plating and imaged by EVOS FL microscope using a 2× objective.

### Immunoblotting

Ara-C-treated cells were lysed in RIPA buffer supplemented with protease and phosphatase inhibitors, and 25μg of total protein were resolved by SDS-PAGE, blotted onto PVDF membranes (Millipore), probed by standard methods, and signals detected by chemoluminescence as described in Supplementary Methods. Primary antibodies are listed in Supplementary Table S1. For preparation of chromatin-bound proteins and soluble nuclear extracts, treated cells were trypsizined, washed in ice-cold PBS, and 2×10^6^ cells were processed using Subcellular Protein Fractionation Kit for cultured cells (Pierce) according to manufacturer’s instructions, and analyzed by Western blotting.

### Cell cycle analysis and intracellular flow cytometry

Cell cycle was analyzed based on DNA content by 4′,6-diamidino-2-phenylindole (DAPI) staining as described in Supplementary methods. Primary antibodies are listed in Supplementary Table S2.

### Replication restart by dual labeling with EdU and BrdU

Cells in culture were pulsed with EdU (10μM) (C10643, Invitrogen) for 30 mins. Plates were then washed and treated with 10μM of Ara-C for 12 hours, washed and released into growth media for the indicated periods of time. BrdU (10μM) (B23151, Invitrogen) was added 30 minutes before harvesting. Trypsinized cells were fixed with 70% ethanol at 4°C overnight, incubated in 2M HCl solution for 30 minutes at RT, and washed in 3% BSA in PBS. EdU was detected using Click-iT EdU Alexa647 picolyl-azide toolkit (C10643, Invitrogen) according to the manufacturer’s protocol. Next, cells were incubated with mouse anti-BrdU monoclonal antibody conjugated to Alexa Fluor 488 (1:20, B35130, Invitrogen) in antibody staining solution (1% BSA, 0.2% Tween-20 in PBS) for 1 hour at RT. After washing in PBS, cells were incubated with DAPI and analyzed by flow cytometry.

### Comet Assay

Alkaline comet assays were performed using CometSlides and Comet Assay kit (Trevigen). See Supplementary methods.

### Immunofluorescence analysis

U2OS cells were seeded 2.5×10^4^ cells/well on 12mm coverslips in 12-well plates, allowed to attach overnight, and treated as indicated. Cells were fixed in 2% paraformaldehyde in PBS for 30 minutes at RT, washed with PBS, permeabilized by 0.5% Triton X-100 in PBS for 10 minutes at RT, and washed with 1mM glycine to quench unreacted aldehydes. Cells were incubated in blocking buffer (0.1% BSA, 0.05% Tween-20, 0.2% Triton X-100, 10% donkey serum) for 1 hour at RT. Coverslips were incubated with primary antibodies specific to γH2A.X, PCNA, pRPA (see Supplementary Table S3) at manufacturer recommended dilutions in blocking buffer overnight at 4°C. After washing in PBS, cells were incubated with AlexaFluor488 or AlexaFluor594-conjugated donkey anti-mouse or anti-rabbit secondary antibodies (Invitrogen) (1:1000) in blocking buffer for 1 hour at RT and washed twice in PBS. Coverslips were mounted using ProLong Gold medium with DAPI (Invitrogen). Slides were examined using a spinning disc confocal (Yokogawa CSU-X1) attached to an inverted microscope (Nikon Eclipse Ti2-E) using a 60× objective (Nikon Plan Apo, 1.4 NA). For quantitative analyses, at least 50 nuclei were selected at random by thresholding based on the DAPI channel, and background-corrected integrated fluorescence density was calculated using ImageJ software. For **BrdU and EdU immunofluorescence analysis**, and **analysis of micronucleation and nuclear morphology** see Supplementary methods.

### Metaphase spreads

U2OS cells were treated with 10μM Ara-C for 12 hours, washed in PBS and released in complete media for 16 hours. The cells were incubated in 50ng/ml colcemid (KaryoMAX, GIBCO) for 12 hours, harvested and swelled with a hypotonic solution (75mM KCl) for 15 minutes at 37°C. The cells were then fixed in ice-cold 3:1 mixture of methanol/acetic acid and dropped on slides. Slides were stained with 5% Giemsa (Sigma), rinsed with distilled water, coverslipped using Permount (Fisher), and scanned on a Keyence BZ-X800 microscope with a 40× NA 0.95 objective.

### RNA-Seq

RNA integrity and purity were assessed by Qubit and Agilent 2100 Bioanalyzer (Agilent Technologies, Santa Clara, CA), and high-quality samples with RIN > 8.0 were sent to Novogene Co. Ltd for library preparation and sequencing on an Illumina NovaSeq 6000 using paired-end 150 chemistry. Sequencing data QC, alignment, quantification, differential expression, and pathway analysis were performed by standard methods as described in Supplementary methods.

### Data sharing

Raw and processed RNA-seq data are available at GEO under the accession code GSE153871.

### Statistical analysis and rigor

Groups were compared using Student’s *t*-test for normally distributed data and Mann-Whitney rank sum test for non-normally distributed data, after testing for normality and equal variance; *p*-values ≤0.05 were considered significant. Where appropriate, one-way ANOVA with Tukey’s HSD *post-hoc* test or two-way ANOVA with *post-hoc* Šídák correction for multiple hypothesis testing were performed. Data were plotted as mean ± standard deviation (SD) unless indicated otherwise. For categorical data, two-tailed Fisher’s exact test was applied. GraphPad Prism software version 7 or higher was used. Identity of cell lines was validated by STR profiling; cultures were routinely tested for Mycoplasma contamination. Image scoring was done blindly; investigators were unblinded after data were collected. *In vivo* studies were repeated twice with comparable results. All other experiments were independently performed three or more times for reproducibility.

## RESULTS

### Cells with *DNMT3A* R882 mutations are more sensitive to replication-stalling drugs

Our prior studies found that *DNMT3A*-mutant cells were relatively resistant to anthracyclines, and showed no differential sensitivity to DNA cross-linking and DSB inducing agents^13^. To investigate the role of DNMT3A in replication stress sensitivity, we treated a panel of leukemia cell lines with and without *DNMT3A*^*R882*^ mutations with the cytosine analog cytarabine and purine analog fludarabine (Fig. 1A,B), purine analog cladribine, and hydroxyurea (HU) which stalls replication by depleting deoxyribonucleotides (Supplementary Fig. S1A,B). Cell lines with *DNMT3A*^*R882*^ SET-2 and KO-52 were more sensitive to replication-stalling drugs than *DNMT3A*^*WT*^ cells K-562 and KU-812, reflected by increased apoptosis measured by annexin V and the cell fraction with sub-G_1_ DNA content (Supplementary Fig. S1C,D).

**Figure 1.**
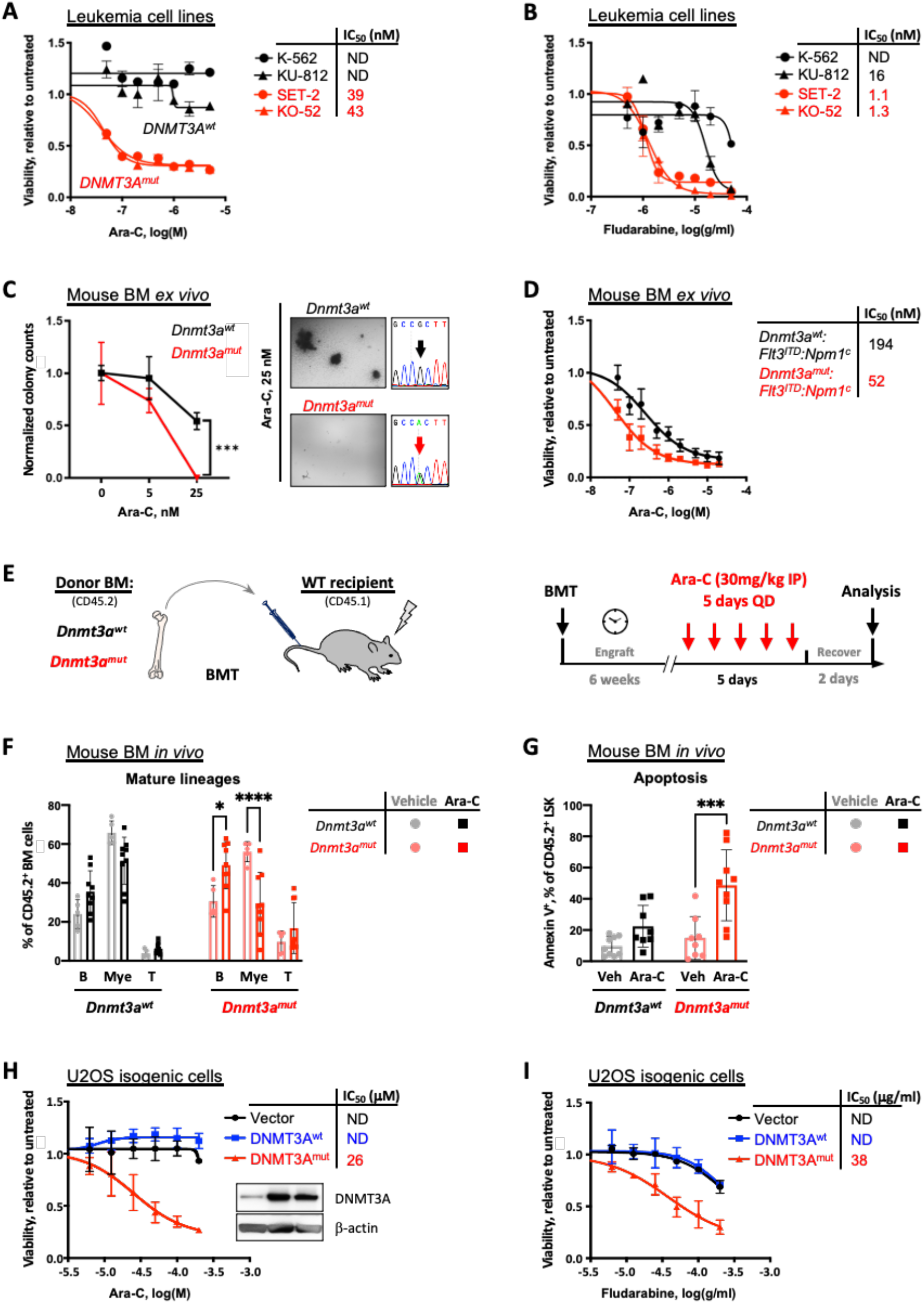
Expression of mutant DNMT3A confers sensitivity to replication stalling agents. (A, B) Sensitivity to elongation-terminating nucleoside analogs cytarabine (Ara-C, A) and fludarabine (B) and corresponding IC_50_ values in leukemia cell lines with *DNMT3A*^*R882*^ (red, SET-2 and KO-52) and *DNMT3A*^*WT*^ (black, K-562 and KU-812) after 48h of treatment, relative to vehicle control, as determined by CellTiter GLO assay in triplicate. (C) Colony formation assay using bone marrow cells derived from *Dnmt3a*^*R878H*^ and littermate *Dnmt3a*^*wt*^ control mice, treated with indicated concentrations of Ara-C and grown in MethoCult M3434, in triplicate. Colonies were scored 10 days after plating and counts normalized to vehicle controls (C). Bone marrow from *Dnmt3a*^*mut*^ mice formed fewer to no colonies compared to wildtype control when treated with 25 nM cytarabine; cDNA Sanger sequencing traces show complete Cre-mediated excision resulting in equal expression of *wt* and *R878H* alleles. (D) *Ex vivo* cytarabine dose responses and corresponding IC_50_ of mouse leukemias driven by *Flt3*^*ITD*^:*Npm1*^*c*^, with and without *Dnmt3a*^*mut*^, as measured by CellTiter GLO assay in triplicate. (E-G) Sensitivity to cytarabine *in vivo*. Congenic CD45.1 recipients reconstituted with *Dnmt3a*^*mut*^ or wild-type control bone marrow cells received 5 daily doses of 30mg/kg Ara-C and were analyzed 48 hours later (E). Frequencies of donor-derived (CD45.2^+^) mature lineage cells (myeloid, B-cells, T-cells) (F) and apoptosis (Annexin V positivity) of the LSK cells (G) in the bone marrow 48 hours after last treatment dose (*n*=5-9, * *p*<0.05, *** *p*<0.001, **** *p*<0.0001, two-way ANOVA with Šídák’s *post-hoc* multiple comparisons test). (H, I) Sensitivity to cytarabine (H) and fludarabine (I) and corresponding IC_50_ values in U2OS cells ectopically expressing wild-type (blue) and R882C mutant (red) forms of *DNMT3A* or empty vector control (black), relative to untreated after 48h of treatment, by AlamarBlue assay. Inset in (H) shows levels of expression of wild-type and mutant DNMT3A, and endogenous DNMT3A in cells transduced with an empty vector control.

We extended these observations to normal and leukemic murine hematopoiesis. Freshly isolated bone marrow cells from mice with conditional *Dnmt3a*^*R878H*^ expression (corresponding to human *DNMT3A*^*R882H*^) plated in the presence of increasing Ara-C concentrations demonstrated a dampened clonogenic potential *ex vivo* in semisolid media compared to wild-type controls (Fig. 1C). Additionally, in a mouse model of AML driven by *Flt3*^*ITD*^ and *Npm1*^*c*^, bone marrow cells from leukemic animals with *Dnmt3a*^*R878H*^ were more sensitive to cytarabine than *Dnmt3a*^*WT*^ in liquid culture *ex vivo* (Fig. 1D). In mice reconstituted with *Dnmt3a*^*R878H*^ but not WT control bone marrow, administration of cytarabine over 5 days *in vivo* led to significant depletion of peripheral blood myeloid (CD11b^+^) cells and a compensatory increase in B-cells (B220^+^), along with elevated apoptosis of hematopoietic stem and progenitor-enriched LSK cell population (Fig. 1E-G). Finally, in primary bone marrow samples from AML patients harboring *FLT3*^*ITD*^ and *NPM1*^*c*^ mutations, presence of *DNMT3A*^*R882*^ was associated with increased *ex vivo* sensitivity to Ara-C compared to *DNMT3A*^*WT*^ (Supplementary Fig. S1E). Together, these data indicate that expression of mutant *DNMT3A* in hematopoietic cells is associated with increased sensitivity to pharmacologically-induced replication stress.

### Cells expressing mutant *DNMT3A* accumulate unresolved DNA damage after cytarabine treatment

To investigate the molecular mechanism of sensitivity to replication-stalling drugs, we lentivirally transduced wild-type and R882C-mutant forms of *DNMT3A* into U2OS cells, a well-established model for DNA damage and repair studies, and confirmed differential cell killing by cytarabine and fludarabine (Fig. 1H,I, and Supplementary Fig. S1F,G). At the same time, steady-state U2OS cell growth was unaffected (Supplementary Fig. S1H).

We next investigated the DNA damage response to cytarabine-induced (approx. IC_20_) replication fork stalling in cells with *DNMT3A*^*R882C*^. Cells overexpressing mutant DNMT3A showed CHK1 phosphorylation persisting over 24 hours of continuous drug exposure and a concomitant increase in p53 phosphorylation, while in *DNMT3A*^*WT*^-expressing cells there was a rapid decline in pCHK1 and a less pronounced p53 activation (Fig. 2A). The intensified signaling in *DNMT3A*^*mut*^-expressing cells reflected accumulation of DNA damage as measured by an increased fraction of mobile DNA per nucleus in an alkaline comet assay (Fig. 2B,C) and confirmed by immunofluorescent detection of phosphorylated histone H2A.X (γH2A.X) (Supplementary Fig. S2A,B). Consistently, SET-2 and KO-52 cell lines with *DNMT3A*^*R882*^ demonstrated accumulation of γH2A.X and cleaved PARP indicative of apoptosis along with persisting low-level pCHK1, while in *DNMT3A*^*WT*^ K-562 and KU-812 cells CHK1 activation was rapidly resolved

**Figure 2.**
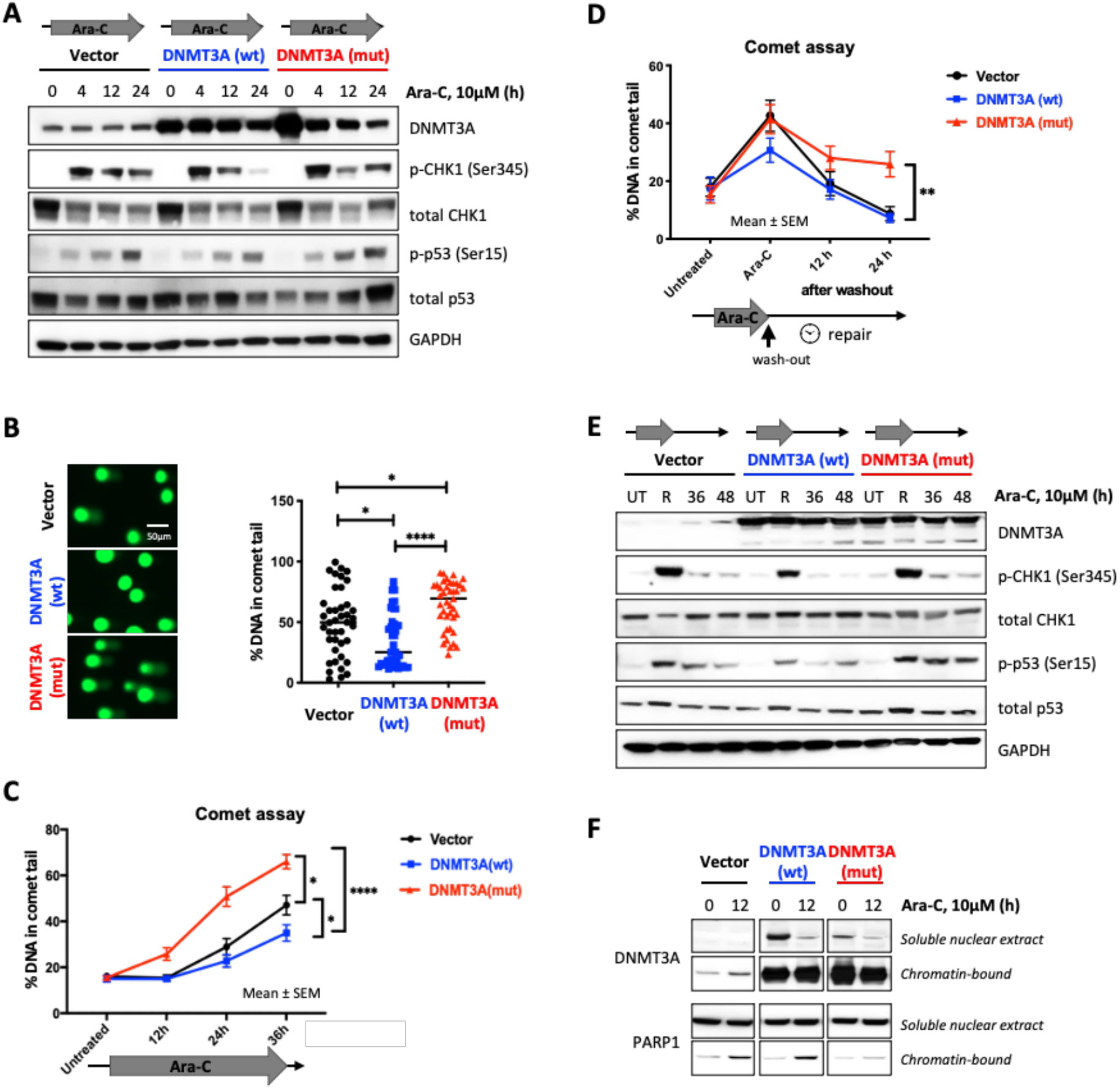
Cells expressing mutant DNMT3A accumulate DNA damage due to incomplete DNA repair upon cytarabine treatment. (A) Immunoblot analysis of the DNA damage signaling at indicated timepoints after cytarabine treatment in U2OS cells with wild-type and mutant *DNMT3A* or empty vector control. (B, C) Detection of damaged DNA by alkaline Comet assay at 24 hours (B) or at indicated timepoints of continuous cytarabine exposure (C) in U2OS cells lentivirally expressing wild-type (blue) and mutant (red) *DNMT3A* or empty vector control (black).(D) Incomplete DNA repair in cells expressing mutant DNMT3A 12 and 24 hours after cytarabine removal following 12h treatment. At least 40 comets per condition were scored using OpenComet plugin for ImageJ software (NIH) by calculating percentage of DNA in the comet tail on the basis of comet head and tail integral intensity. (* *p*<0.05, ** *p*<0.01, **** *p*<0.0001, Mann-Whitney rank-sum test; graphs represent mean ± SEM). (E) Analysis of DNA damage response in U2OS cells lentivirally expressing wild-type and mutant DNMT3A or empty vector control at 12 hours of cytarabine exposure and 36 and 48 hours after drug wash-out (UT, untreated; R, at release). (F) Recruitment of PARP1 to chromatin from soluble nuclear extract after cytarabine treatment in cells overexpressing wild-type or mutant forms of *DNMT3A*.

(Supplementary Fig. S2C). These results indicate that cells with mutant *DNMT3A* accumulate DNA damage after cytarabine.

Deficiency in DNA repair is a common explanation for accumulation of DNA damage in drug treated cells. Accordingly, DNA damage resolution after cytarabine wash-out was markedly delayed in *DNMT3A*^*R882C*^- overexpressing cells and coincided with sustained p53 activation (Fig. 2D,E). Of note, per-cell levels of a DNA damage marker γH2A.X were also significantly higher (Supplementary Fig. S2D,E) consistent with a defect in DNA damage repair. Mechanistically, this coincided with impaired recruitment of PARP1, a key component in directing DNA repair^42^, to chromatin in *DNMT3A*^*mut*^-expressing cells (Fig. 2F).

### Cells expressing mutant *DNMT3A* are prone to replication stress and fork collapse

We hypothesized that increased sensitivity to DNA damage during S-phase and accumulation of unresolved DNA breaks was associated with replication stress. Indeed, this was found in cells expressing mutant *DNMT3A* as evidenced by high levels of phospho-RPA, which persisted even after drug wash-out (Fig. 3A,B), and coincided with γH2A.X accumulation (Supplementary Fig. S3A,B). Consistently, *DNMT3A*^*mut*^- expressing cells treated with cytarabine were characterized by a higher number and a larger area of PCNA foci^43^ (Fig. 3C,D). Of note, since Ara-C inhibits elongation, BrdU incorporation, an otherwise ideal means to visualize replication foci, could not be used in these studies.

**Figure 3.**
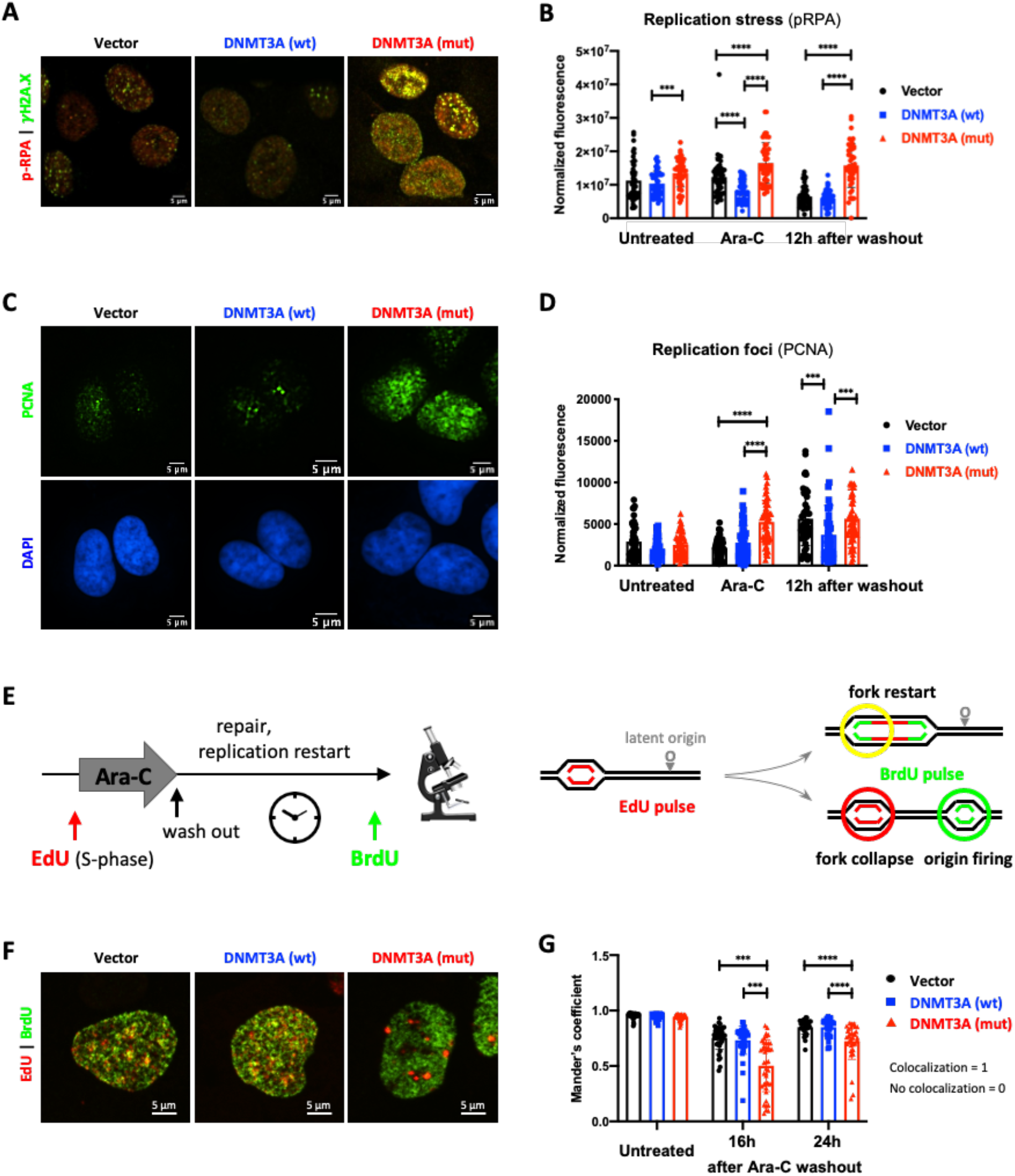
Cells expressing mutant *DNMT3A* experience prolonged replication stress after cytarabine treatment. (A, B) Replication stress and DNA damage after cytarabine treatment in U2OS cells transduced to express wild-type and mutant *DNMT3A* or empty vector control. Representative immunofluorescence microscopy images after 12h exposure (A, pRPA pSer33, red, and DSB marker γH2A.X, green) and pRPA signal quantification at steady state, 12 hours of treatment, and 12 hours after drug wash-out in at least 50 cells per condition (B). (C, D) Analysis of replication foci by PCNA immunofluorescence staining (green) and DAPI (blue) after 12 hours of cytarabine treatment (C) and PCNA signal quantification in untreated, treated, and 12 hours after drug removal in at least 50 cells per condition (D) in U2OS cells with wild-type and mutant *DNMT3A*. (E-G) Replication restart analysis by EdU and BrdU double-labeling pulse-chase experiments. U2OS cells overexpressing wild-type and mutant *DNMT3A* or empty vector control were pulsed with EdU (red) for 1 hour, treated with cytarabine for 12 hours, washed and pulsed with BrdU (green) at indicated timepoints (E). Representative immunofluorescence microscopy images of double-labeled cells (F, 16 hours after drug removal) and quantification of BrdU and EdU signal colocalization (Manders coefficient) in at least 25 nuclei per condition at indicated time points after drug removal (G). (*** *p*<0.001, **** *p*<0.0001, Mann-Whitney rank sum test).

Next, we examined resumption of DNA synthesis using pulse-chase double-labeling experiments with EdU (to identify replicating cells) and, following cytarabine wash-out, BrdU (marking replication restart). Cells were able to resume replication regardless of the *DNMT3A* genotype, with similar overall kinetics (Supplementary Fig. S3C,D) despite persisting DNA damage at replication forks. Confocal imaging revealed that *DNMT3A* mutant cells were more likely to suffer fork collapse seen as lack of colocalization between EdU (replication foci pre-cytarabine) and BrdU (replication restart) (Fig. 3E-G).

### Cells expressing mutant *DNMT3A* progress through the cell cycle despite persisting DNA damage, leading to mitotic catastrophe

The ability of cytarabine-treated U2OS cells expressing wild-type or mutant *DNMT3A* to resume cycle progression was determined. The analysis was facilitated by partial cell synchronization at the G_1_/S phase boundary by Ara-C treatment. By 18h after drug wash-out, cells completed replication and began entering mitosis marked by histone H3 phosphorylation (Fig. 4A and Supplementary Fig. S4A,B). Notably, cells with *DNMT3A* mutation continued to advance through the cell cycle, albeit with a lag (Supplementary Fig. S4C), despite unresolved DNA damage. As the result, there was an increased frequency of chromosome breaks in metaphase preparations of mutant DNMT3A-expressing cells (Fig. 4B,C) along with hallmarks of mitotic catastrophe such as micronucleation and abnormal nuclear morphology (Fig. 4D-G).

**Figure 4.**
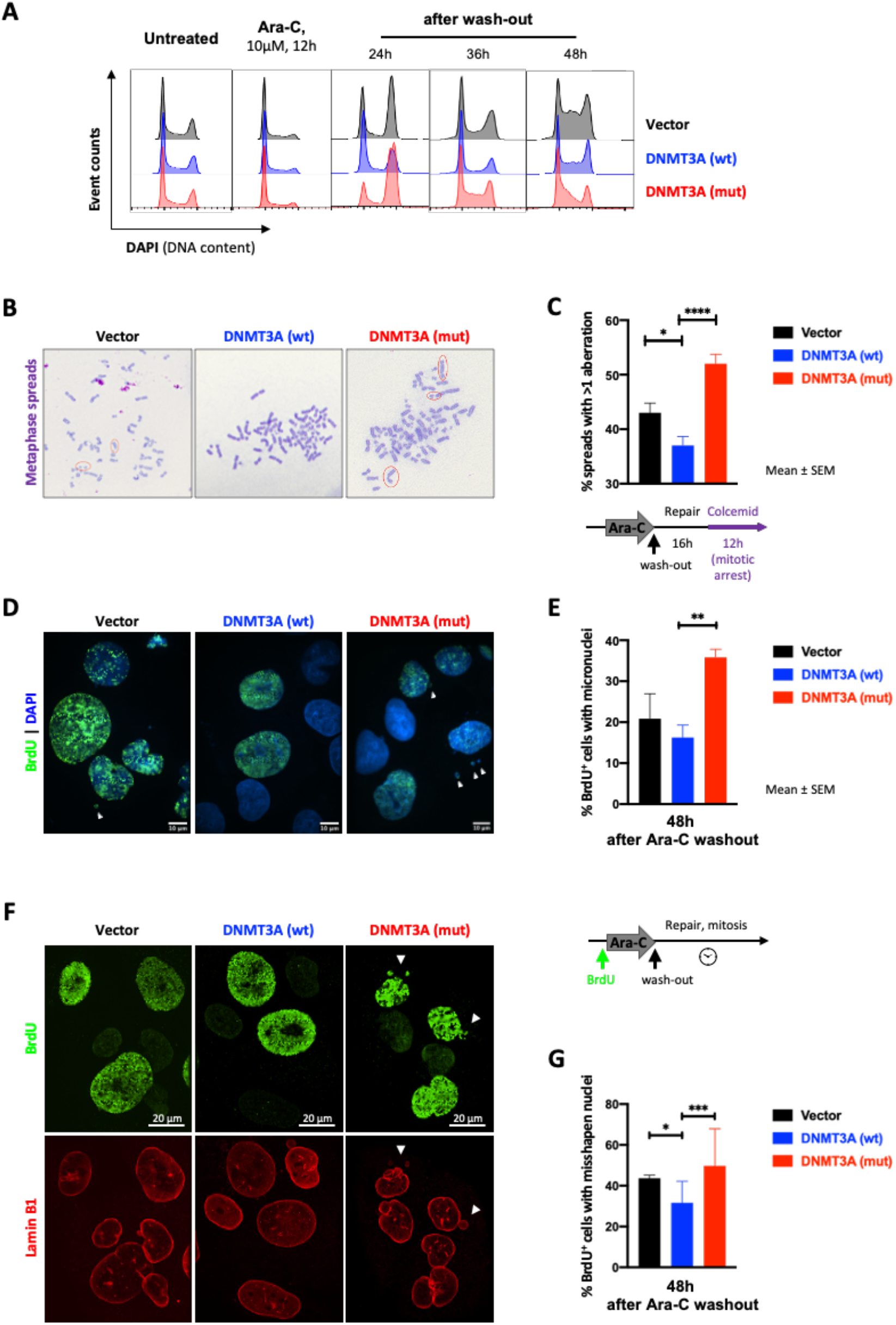
Mutant DNMT3A-expressing cells carrying DNA breaks after cytarabine treatment progress through mitosis. (A) Cell cycle profiles (by DNA content) in U2OS cells with wild-type (blue) and mutant (red) *DNMT3A* or empty vector control (black) after cytarabine treatment and after wash-out. Fixed and permeabilized cells were stained with DAPI and analyzed by flow cytometry. (B, C) Persistent DNA breaks in mitosis after cytarabine treatment in U2OS cells lentivirally overexpressing wild-type and mutant *DNMT3A* or empty vector control. Representative images of metaphase spreads (B, Wright-Geimsa stain) and frequency of metaphases with chromosomal abnormalities from five independent experiments each scoring at least 200 metaphases per genotype (C) 16 hours after drug wash-out (* *p*<0.05, **** *p*<0.0001, unpaired *t*-test; graphs represent mean ± SEM). (D-G) Mitotic catastrophe as a consequence of abnormal mitoses in cells with persistent DNA damage after cytarabine wash-out. Cells were pulsed with BrdU for 1h, exposed to cytarabine for 24h, and released from drug treatment; 48 hours later cells were fixed and stained for BrdU (D, E) and Lamin-B1 (F, G). Representative microphotographs of nuclear morphology (D, DAPI was used to visualize all nuclei in blue) and percentage of BrdU^+^ cells with micronuclei (E, ** *p*<0.01, unpaired *t*-test; graphs represent mean ± SEM from three independent replicate experiments). Representative microphotographs of nuclear lamina abnormalities visualized by Lamin-B1 staining (red) in BrdU^+^ cells (green) (F) and fraction of BrdU^+^ cells with abnormally-shaped nuclei (G). At least 100 nuclei per genotype were scored in each of two independent experiments (* *p*<0.05, *** *p*<0.001, pair-wise two-tailed Fisher’s exact tests, where each normal nucleus was categorized as 0 and each abnormal as 1).

Collectively, our studies indicate that cells with mutant DNMT3A exhibit a defect in repair and recovery of stalled replication forks, resulting in accumulation of DNA breaks carried through mitosis. The accentuated cytarabine sensitivity is further highlighted by our observation that doubling Ara-C concentration prevented mutant DNMT3A-expressing cells from completing replication seen as G_2_ peak degradation in cell cycle analysis, together with a new sub-G_1_ population indicative of cell death, in contrast to DNMT3A wild-type cells that maintained a normal cell cycle distribution (Supplementary Fig. S4D).

### Gene expression profiling after cytarabine treatment identifies signatures of deregulated cell cycle and DNA replication in cells expressing mutant DNMT3A

To gain a broader view of cytarabine response, we performed RNA-sequencing to elucidate gene expression signatures deregulated by mutant DNMT3A (Supplementary Table S4). Unsupervised hierarchical clustering of the 2000 most variable genes demonstrated predominant gene activation induced by cytarabine treatment and robust separation from untreated controls, with consistency between biological triplicates (Supplementary Fig. S5A). Pairwise comparisons between cytarabine-treated and untreated control cells found wide-spread gene activation that was largely shared between genotypes yet featured a significant proportion of uniquely regulated genes in *DNMT3A*^*mut*^-expressing samples (Fig. 5A and Supplementary Fig. S5B; Supplementary Table S5).

**Figure 5.**
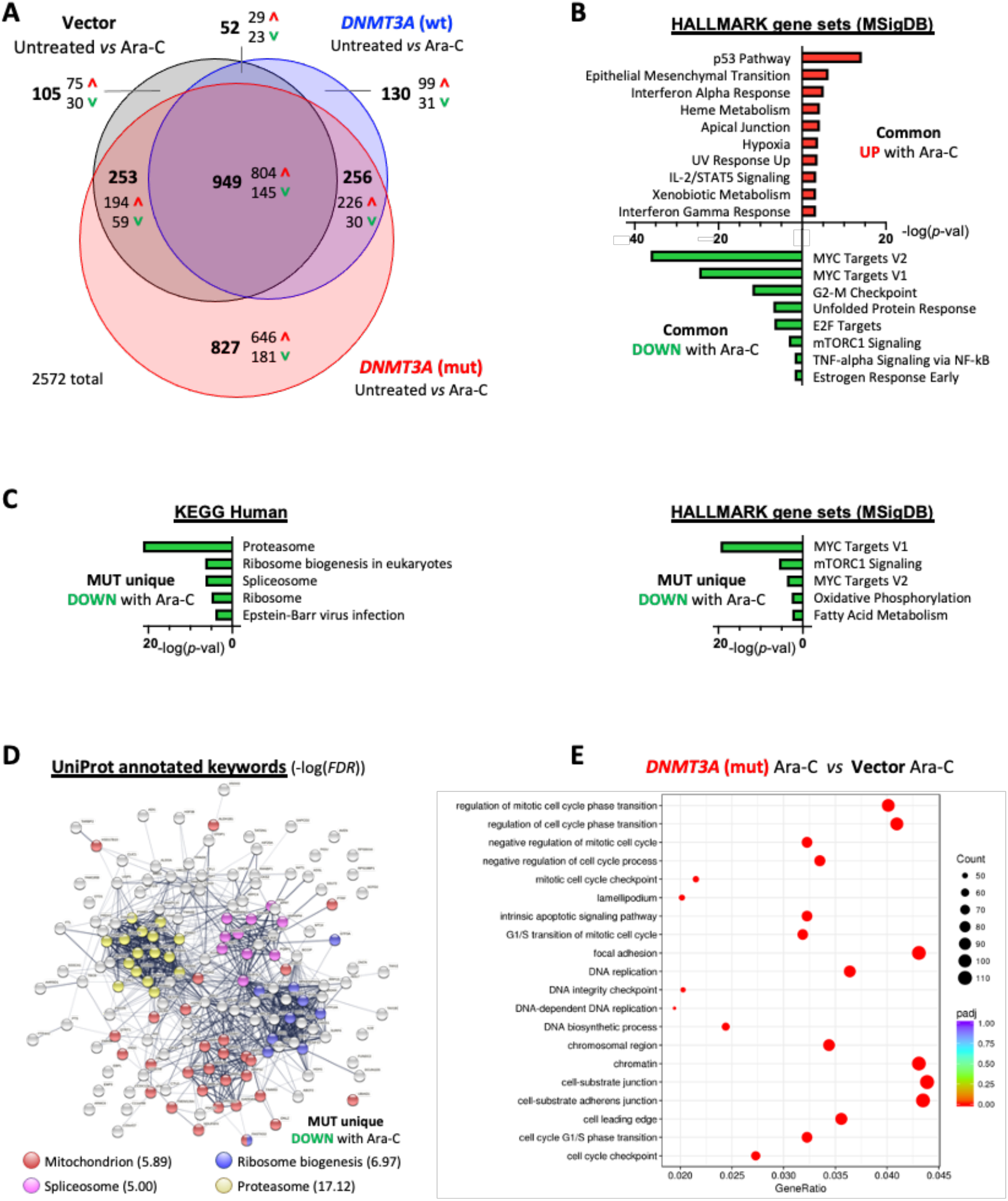
Gene expression profiling identifies pathways deregulated in mutant DNMT3A expressing cells treated with cytarabine. (A) Venn diagram showing differentially expressed genes (FC:21.5, FDR<0.05) between vehicle and Ara-C-treated (10 μM, 24h) cells that are common or unique to each of the experimental groups: U2OS cells expressing wild-type (blue) or mutant (red) *DNMT3A*, or empty vector control (black). (B) HALLMARK gene sets (MSigDB) significantly enriched among commonly up- or downregulated genes in all three groups (FDR<0.1). (C) Significantly enriched KEGG pathways and HALLMARK gene sets among genes uniquely downregulated in *DNMT3A*^*mut*^ cells after cytarabine treatment (FDR<0.1). (D) STRING functional protein network analysis corresponding to genes uniquely downregulated in the *DNMT3A*^*mut*^ cells 24 hours after treatment with Ara-C, and their UniProt annotations. (E) Enrichment of GO terms among genes differentially expressed (*p*<0.05) in cytarabine-treated *DNMT3A*^*mut*^ cells compared to treated empty vector controls.

Pathway analysis using Enrichr to search against HALLMARK gene set collection (MSigDB) and Kyoto Encyclopedia of Genes and Genomes (KEGG) database (Supplementary Table S6) identified p53 signaling as most significantly overrepresented among commonly upregulated genes in all groups (Fig. 5B), consistent with genotoxic stress. At the same time, genes associated with the MYC pathway, G2/M checkpoint, and E2F targets were repressed in all groups, indicative of ongoing replication stress and proliferation delay, reinforcing our biochemical findings (Fig. 2A,E and 4A). Conversely, genes uniquely upregulated in *DNMT3A*^*mut*^-expressing cells after cytarabine treatment did not show similarly robust gene set enrichment apart from high expression of genes implicated in stemness and hematopoietic development such as *CD34*^*44*^, *PDGFRA*^*45*^, *HOXB6*^*46*^, *IKZF2*^*47*^ (Supplementary Fig. S5B,C). Analysis of genes with decreased expression after cytarabine uniquely in the mutant DNMT3A-expressing cells highlighted further repression of proliferation-related MYC targets and identified disruption of vital leukemia and stem cell regulatory and metabolic pathways such as mitochondrial oxidative phosphorylation (OXPHOS) and fatty acid metabolism, RNA-splicing, and protein synthesis and degradation^48-51^ (Fig. 5C). Furthermore, functional networks formed by proteins encoded by these exclusively DNMT3A^mut^-downregulated genes were characterized by prominent clusters annotated as proteasome, mitochondria, spliceosome, and ribosome biogenesis (Fig. 5D), suggesting distinctive shifts in cell physiology and molecular liabilities.

Pairwise comparison between cytarabine-treated DNMT3A-mutant and empty vector control cells found deregulation of cell cycle-related pathways, emphasizing G1/S phase transition and DNA replication as well as cell adhesion^52^ (Fig. 5E). Gene set enrichment analysis (GSEA) of gene ontology (GO) datasets additionally detected negative enrichment of signatures related to chromatin packaging and myeloid cell differentiation, in line with previous reports and a prominent role of *DNMT3A* in myeloid biology^5,11,13,24,53^ (Supplementary Fig. S5D).

## DISCUSSION

Mutations in *DNMT3A* are among the most common genetic alterations in *de novo* AML and are associated with adverse outcomes^6-9,12,14,15,22,54,55^. In particular, the R882 mutation conveys an unfavorable prognosis in patients ≥60 years of age^56^. This is at least in part due to relative resistance to anthracyclines stemming from another defect in DNA damage response^12-14^ and inability to tolerate dose-sense regimens, straining an already limited list of therapeutic options available for this patient group^3,19^. Here we show that cells with *DNMT3A* mutations may be sensitive to replication stress-related DNA damage induced by nucleoside analogs such as cytarabine, in agreement with a recent clinical trial^21^. Low-dose cytarabine-, cladribine-based, or other similar regimens are well-tolerated and hence suitable for older patients and those with poor performance status.

A growing body of evidence recognizes emergence of “collateral sensitivities”, or increased susceptibility to a second therapy as a fitness trade-off in cancer cells following the development of resistance to initial treatment or other perturbations^57-59^. This raises a possibility of rationally constructing synthetic lethality-based treatment approaches exploiting such antagonistic pleiotropic mechanisms^60,61^. The concept that the same genetic defect may be protective in the context of a specific type of DNA damage yet make cells vulnerable to a different type of genotoxic insult has been described^62^. Similarly, our studies demonstrate whereas the presence of *DNMT3A*(R882) decreases the sensitivity to DNA torsional stress induced by anthracyclines and possibly other DNA intercalators^13^, it renders the cells more susceptible to the deleterious effects of replication fork stalling.

Accentuated replication stress in cells expressing mutant DNMT3A is further supported by previous findings of negatively enriched E2F target and CHEK1-regulated G_2_/M checkpoint gene sets reported in independent AML cohorts as well as in mice^13,28^, which reflect G_1_/S cell cycle phase transition rate^29,30,63^ and G_2_/M checkpoint adaptation^64,65^. Accordingly, after low-dose cytarabine exposure (approx. IC_20_) mutant DNMT3A-expressing cells demonstrated delayed cell cycle progression that culminated in entering mitosis with damaged DNA. Surviving cells demonstrated increased formation of micronuclei, structures containing shattered chromosomes, indicative of mitotic catastrophe^41,66^. Furthermore, incremental increase in cytarabine concentration resulted in disproportionate apoptotic response, highlighting heightened replication stress in *DNMT3A*-mutant cells as a potential therapeutic vulnerability that can be unveiled by replication-stalling nucleoside analogs.

Such distinctive sensitivity to cytarabine arises following a defect in recovery from replication fork arrest that leads to accumulation of persistent, unrepaired DNA damage observed in cells with mutant DNMT3A. While previous studies have attributed specific DNA damage repair pathways as defining therapeutic responses to replication stress-inducing nucleoside analogs^33,34,67-70^, the exact mechanism(s) that underlie impaired resolution of cytarabine-induced DNA lesions in cells with mutant DNMT3A remain uncertain. Interestingly, other than a slight upregulation of genes annotated to be repressed during UV response indicative of a possible disruption of nucleotide excision repair^42^ (Supplementary Fig. S5C) our gene expression studies did not detect signatures of impaired DNA damage repair. Prior in-depth studies reported only modest changes in DNA methylation^13,25-27,71^ that predominantly affected lineage-specific regulatory elements^28,53,72-74^ yet did not affect DNA damage and repair pathways. Whether a methylation-independent mechanism of mutant DNMT3A action might be at play remains to be determined. In support of this, wild-type DNMT3A protein was found to directly associate with stalled replication forks^31^ and likely enriched in the PCNA interactome^75^. There are many examples of non-catalytic accessory functions of epigenetic enzymes regulating both histone^76,77^ and DNA modifications^78-80^. Future studies interrogating the replication fork-associated proteome^81^ in cells with and without mutant DNMT3A in addition to differential recruitment of DNA repair proteins to chromatin and proximity-labeling proteomics approaches^82,83^ will be necessary to clarify its role in preserving DNA integrity. This emerging knowledge will be instructive to develop further therapeutic combination strategies.

In addition to DNA replication, cell cycle regulation, and proliferation-related gene sets, which are likely secondary to DNA damage after cytarabine, our gene expression studies uncovered signatures of deregulated cell adhesion, proteostasis, as well as repressed OXPHOS and fatty acid metabolism, and RNA splicing, all of which are critical for hematopoietic stem cell function and are commonly deregulated during aging and in myeloid malignancies. Notably, deregulation of ribosome biogenesis and protein degradation may reflect loss of stemness^84,85^, whereas increased reliance on OXPHOS and fatty acid metabolism are characteristic of chemoresistant acute myeloid leukemia^49,52,86^ and together with altered splicing engender potential therapeutic vulnerabilities^50,87,88^. It will be interesting to investigate if these processes contribute to the leukemic progression in individuals with clonal hematopoiesis with *DNMT3A* mutations^89-94^ to serve as biomarkers for high risk of AML development.

In summary, our studies demonstrate that cells expressing mutant DNMT3A have a defect in recovery from replication fork arrest and subsequent accumulation of unresolved DNA damage, which may have therapeutic tractability. These results demonstrate, in addition to its role in epigenetic control, DNMT3A contributes to preserving genome integrity during DNA replication and suggest that cytarabine-induced replication fork stalling may further synergize with other agents aimed at DNA damage and repair, replication, or cell survival, as well as splicing and oxidative phosphorylation^50,51,87,88,95-97^. Future studies will determine if such combinatorial treatment approaches can be developed to usher in the era of precision oncology for AML patients with mutant *DNMT3A*.

## ACKNOWLEDGEMENTS

The authors gratefully acknowledge generous support by NIH awards CA178191 and DK121831 (O.A.G.), CA188561 (R.O.), and CA252400 (B.K.L.). O.A.G. was supported in part by the Ocala Royal Dames for Cancer Research, the Harry T. Mangurian Foundation, and the Thomas H. Maren Junior Investigator Fund. B.K.L. was supported in part by the Florida Breast Cancer Foundation and the Ocala Royal Dames for Cancer Research. J.L. and D.D.R. are Leukemia and Lymphoma Society (LLS) Special Fellows; J.D.L. is supported in part by the LLS Specialized Center of Research (SCOR). Immunophenotyping was performed at the Flow Cytometry and Imaging Core, UF Interdisciplinary Center for Biotechnology Research (ICBR) (RRID: SCR_019119).

## AUTHORSHIP CONTRIBUTIONS

K.V. and O.A.G conceived the study and designed experiments, with conceptual input from J.D.L., R.O., C.R.C., and B.K.L.

K.V. performed experiments, with assistance from Y.F., D.E.S., and O.A.G., and additional help from C.M.B., K.I.K., C.T., Z.Z., H.L.C.R., L.M.P., C.G., J.L., D.D.-R., and R.L.B.

S.K. and P.B.S. performed and analyzed *ex vivo* drug studies in primary AML samples.

P.N. analyzed RNA-seq dataset, with assistance from A.R. and S.P.

J.E.B. assisted with microscopy experiments and data analysis.

K.V. and O.A.G. analyzed the data and wrote the manuscript, with input from all co-authors.

## CONFLICT OF INTEREST DISCLOSURES

The authors declare no relevant competing financial interests.

## SUPPLEMENTARY INFORMATION

## Supplementary methods

### Animal studies

A conditional *Dnmt3a*^*R878H*^ knock-in mouse line (Jackson Laboratory stock No. 031514) was previously described^1^. Expression of the mutant form of *Dnmt3a*, also referred to as *Dnmt3a*^*mut*^, is activated by a hematopoietic-specific *Mx1*-Cre deletor following poly(I:C) injections; poly(I:C)-treated *Dnmt3a*^*+/+*^:*Mx1*-Cre^+^ are used as controls. Successful recombination and expression of the point mutant is verified by Sanger sequencing of the peripheral blood nucleated cell cDNA as previously reported.

To generate mouse leukemias with and without *Dnmt3a*^*R878H*^, we intercrossed *Dnmt3a*^*R878H*^:*Mx1-Cre* mice with animals harboring constitutive *Flt3*^*ITD*^ and inducible *Npm1*^*c*^ alleles as previously described. Bone marrow from *Dnmt3a*^*mut*^:*Flt3*^*ITD*^:*Npm1*^*c*^ and control *Dnmt3a*^*wt*^:*Flt3*^*ITD*^:*Npm1*^*c*^ animals developing leukemia after Cre induction (WBC reach or exceed 50×10^3^/μl) is harvested for *ex vivo* drug dose response analyses.

For *in vivo* cytarabine treatment studies, 0.5×10^6^ CD45.2 donor bone marrow cells from fully excised *Dnmt3a*^*+/R878H*^*:Mx1-Cre*^*+*^ (*Dnmt3a*^*mut*^) and littermate control *Dnmt3a*^*+/+*^*:Mx1-Cre*^*+*^ *(Dnmt3a*^*wt*^) mice were injected through tail veins into lethally irradiated congenic CD45.1 recipients and allowed to engraft for 6 weeks. Successfully transplanted animals of both genotypes, confirmed by CD45.1/CD45.2 peripheral blood chimerism, were randomized between treatment (30mg/kg cytarabine/day for 5 days IP) and vehicle (PBS) (*n*=5/group). Bone marrow was harvested 48h after last injection by spin-flush method and red blood cell lysis. Analysis of the bone marrow cellular composition was done by flow cytometry. Apoptosis in the stem/progenitor-enriched Lineage^−^Sca1^+^cKit^+^ (LSK) population was determined by Annexin V (556422, BD Pharmingen) binding and 4′,6-diamidino-2-phenylindole (DAPI) uptake (Thermo Fisher). All cell surface antibodies were from eBioscience or BioLegend: NK1.1 (PK136), CD11b (M1/70), CD45R (RA3-6B2), CD3 (17A2), Gr-1 (RB6-8C5), Ter119 (TER119), CD19 (6D5), CD4 (GK1.5), CD8 (53-6.7), cKit (2B8), Sca-1 (D7), CD150 (TC15-12F12.2), CD48 (HM48-1), CD16/32 (93), CD34 (RAM34), Ki67 (16A8), CD45.1 (A20), CD45.2 (104).

### Primary AML samples

Bone marrow aspirates from patients with AML driven by *FLT3*^*ITD*^ and *NPM1*^*c*^, with (*n*=4) and without (*n*=5) *DNMT3A*^*R882*^, were subjected to *ex vivo* drug dose response study as previously described^1^, approved by the ethical committee of the Medical University of Vienna #EK Nr: 2008/2015.

### Apoptosis assay

Apoptosis was measured by flow cytometry using Annexin V (APC) and propidium iodide (PI) double staining. Cells were treated as indicated, collected, counted, and 1×10^6^ cells were washed with ice-cold PBS twice, and resuspended in 200 μl of 1X binding buffer containing Annexin V and PI according to the manufacturer’s instructions for 15 minutes at 37°C in the dark. The proportion of viable, apoptotic and necrotic cells was quantified by flow cytometry and analyzed by the FlowJo software.

### Cell growth assay by methylene blue staining

Lentivirally transduced U2OS cells were plated 1×10^3^ cells/well in 96-well plates. Every day, triplicate wells were fixed and stained by 0.5% methylene blue (Sigma) in 50% methanol, then washed in water and allowed to air-dry. At experiment completion, bound methylene blue was solubilized in 0.1% SDS, and OD_595_ was measured using the SpectraMax plate reader. The amount of bound methylene blue is proportional to the cell number; relative cell growth was normalized to day 0 set at 1.

### Immunoblotting

Cells treated with Ara-C were collected after washing in ice-cold PBS and lysed in RIPA buffer supplemented with protease and phosphatase inhibitors. Protein concentrations were determined using BCA protein assay (Pierce). For target detection by Western blotting, 25μg of total protein were resolved by SDS-PAGE on 4–12% Bis-Tris gels (Invitrogen), blotted onto PVDF membranes (Millipore) and probed by standard methods. Primary antigen-specific antibodies, listed in Supplementary Table S1, were used at 1:1000 dilutions and signals were detected after incubation with secondary HRP-linked species-specific antibodies (Santa Cruz Biotech, 1:5000 dilution) using KwikQuant chemiluminescence reagents and a digital imaging station (Kindle Biosciences).

### Comet Assay

Alkaline comet assays were performed using CometSlides and Comet Assay kit (Trevigen), according to manufacturer’s instructions. Cells were treated as indicated, harvested, counted, and embedded in low melting point agarose buffer layered onto CometSlides. After processing and electrophoresis, slides were stained with SYBR Gold and imaged using EVOS FL (Life Technologies) imaging system using a 10× NA 0.30 objective. Images were analyzed using ImageJ software (NIH) and an OpenComet plugin calculating percent damaged DNA on the basis of comet head and tail sizes (measured in pixels) and their integral fluorescence intensity.

### Cell cycle analysis

Cell cycle was analyzed based on DNA content by DAPI staining. Cells were treated as indicated, harvested, and counted. Adherent cells were trypsinized as necessary. One million cells were washed in PBS and fixed in 3.7% formaldehyde for 15 minutes at room temperature (RT) and permeabilized in 0.5% Triton X-100 in PBS for 15 minutes. Alternatively, cells were extensively washed and fixed in 70% ice-cold ethanol at 4°C overnight. The cells were then washed in PBS, stained with 1μg/ml DAPI (Thermo Fisher Scientific) in PBS for 10 minutes, and analyzed by flow cytometry using BD LSR Fortessa. Where indicated, cells were incubated with antibodies against phosphorylated histone H3 (pS28) or H2A.X (pS139) conjugated to AlexaFluor647, for 20 minutes. After washing with PBS, DNA was counterstained using 1μg/ml DAPI (Thermo Fisher) in PBS for 10 minutes at RT, and analyzed by flow cytometry. DNA content histograms were generated using FlowJo software version 10.4.1, using the CellCycle plugin as appropriate.

### BrdU and EdU pulse-chase double-labeling experiments

U2OS cells were pulsed with EdU (10μM) (C10340, Invitrogen) for 30 minutes. The cells were then washed and treated with 10μM of Ara-C for 12 hours and then released in BrdU (10μM) for 30 minutes at the indicated time points. Cells were fixed and permeabilized as described above. Cells were incubated in 2M HCl solution for 30 minutes at RT and neutralized in a solution containing 3% BSA in PBS. EdU was detected using the Click-iT EdU AlexaFluor647 imaging kit (C10340, Invitrogen) according to the manufacturer’s protocol. Cells were then incubated in blocking buffer (0.1% BSA, 0.05% Tween 20, 0.2% TritonX-100, 10% donkey serum) for 1 hour at RT followed by 1:200 mouse anti-BrdU monoclonal antibody in blocking buffer overnight at 4°C. After washing in PBS, cells were incubated with AlexaFluor488-labelled donkey anti-mouse secondary antibody (Invitrogen) (1: 1000) in blocking buffer for 1 hour at RT and washed twice in PBS. Coverslips were mounted with ProLong Gold with DAPI (Invitrogen). Slides were examined on Nikon Eclipse Ti2 confocal microscope using a 63× NA 1.46 objective. For quantitative analyses, at least 50 nuclei were selected at random and analyzed using the ImageJ or NiS software. Colocalization was quantified using Manders overlap coefficient, which measures the degree of overlap of two separate fluorescence channels relative to the total intensity within each channel. A Manders coefficient of 0 corresponds to no colocalization, whereas a value of 1 means 100% colocalization.

### Analysis of micronucleation and nuclear morphology

Exponentially growing U2OS cells on glass coverslips were pulsed with BrdU (10μM) for 60 minutes, washed with PBS, treated with 10μM Ara-C for 24 hours, then released into complete media for the indicated duration of time.

Cells were fixed, permeabilized, and extracted in 2M HCl as discussed above. Coverslips were then incubated in blocking buffer followed by anti-BrdU mouse monoclonal antibody and anti-Lamin B1 rabbit polyclonal antibody (ab16048, Abcam) in blocking buffer overnight at 4°C. After washing in PBS, cells were incubated with AlexaFluor488-labelled or AlexaFluor594-conjugated donkey anti-mouse or anti-rabbit secondary antibodies, washed twice in PBS, and mounted in ProLong Gold mounting medium with DAPI. The images were taken on the Nikon Eclipse Ti2 confocal microscope using a 100× NA 1.49 objective. At least 100 randomly selected BrdU-positive nuclei were counted for presence of micronuclei, or analyzed for eccentricity using the ImageJ and the NiS Elements software.

### RNA-seq analysis

RNA integrity and purity were assessed by Qubit and Agilent 2100 Bioanalyzer (Agilent Technologies, Santa Clara, CA), and high-quality samples with RIN > 8.0 were sent to Novogene Co. Ltd for library preparation and sequencing. NEBNext^®^ Ultra™ RNA Library Prep Kit was used for poly-A tailed mRNA enrichment, fragmentation, cDNA synthesis, NEBNext hairpin loop adaptor addition and library preparation. The library quality and insert size were assessed on Agilent 2100 Bioanalyzer and the concentration was determined by Qubit and qPCR methods. The libraries were then sequenced on an Illumina NovaSeq 6000 to generate PE 150 reads.

Raw reads that were pre-processed to remove low quality reads, poly-Ns and adapter sequences were provided in FASTQ format by Novogene and used for all downstream analyses. Clean FASTQ reads were uploaded into PartekFlow^®^ software, version 8.0 (8.0.19.0322 build, Partek, Inc.) on a standalone server at UF ICBR, followed by pre-alignment QA/QC. These reads were then mapped to the human reference genome build 38 (hg38) with the default settings in the STAR aligner version 2.5.3a followed by post-alignment QA/QC showing average alignment rate 93-95% and 56.4 million transcriptome-aligned reads per sample. The transcript read count was obtained using the default strict paired end compatibility and minimum reads of 10 in the Quantify to Annotation Model (Partek E/M) method and quantified to human hg38 – Ensembl Transcripts release 95. About 80% of the reads mapped fully to the exons indicating an overall good read distribution and rRNA depletion. Low expression genes (read counts <10) were filtered to reduce noise and raw read counts across samples were normalized using TMM with an offset of 1 (to avoid zero counts during further logarithmic transformation).

For differential gene expression analysis, first unsupervised hierarchical clustering was performed on log2 transformed TMM values using Pearson correlation clustering with average linkage. Cluster 3.0^2^ was used to log2 transform input values, filter top 2000 most variable genes and cluster the data. Morpheus online tool (https://software.broadinstitute.org/morpheus) was used for visualization. Heatmap used color coding where minimum and maximum values were set for each row individually. Differentially expressed genes between vehicle and cytarabine-treated cells in each genotype were determined by edgeR quasi-likelihood (QL) pipeline^3^ and filtered based on fold change ≥1.5 and FDR <0.05. Overlap between groups and proportional Venn diagrams were constructed by Venny online javascript tool (https://github.com/benfred/venn.js; https://www.stefanjol.nl/venny). Gene ontology (GO) and pathway enrichment analyses were done using Enrichr^4,5^ and GSEA^6,7^ to search against MSigDB Hallmark^8^ and KEGG^9^ databases; enrichment of protein-protein association networks was visualized using STRING^10-12^ application with default parameters.

**Supplementary Figure S1.**
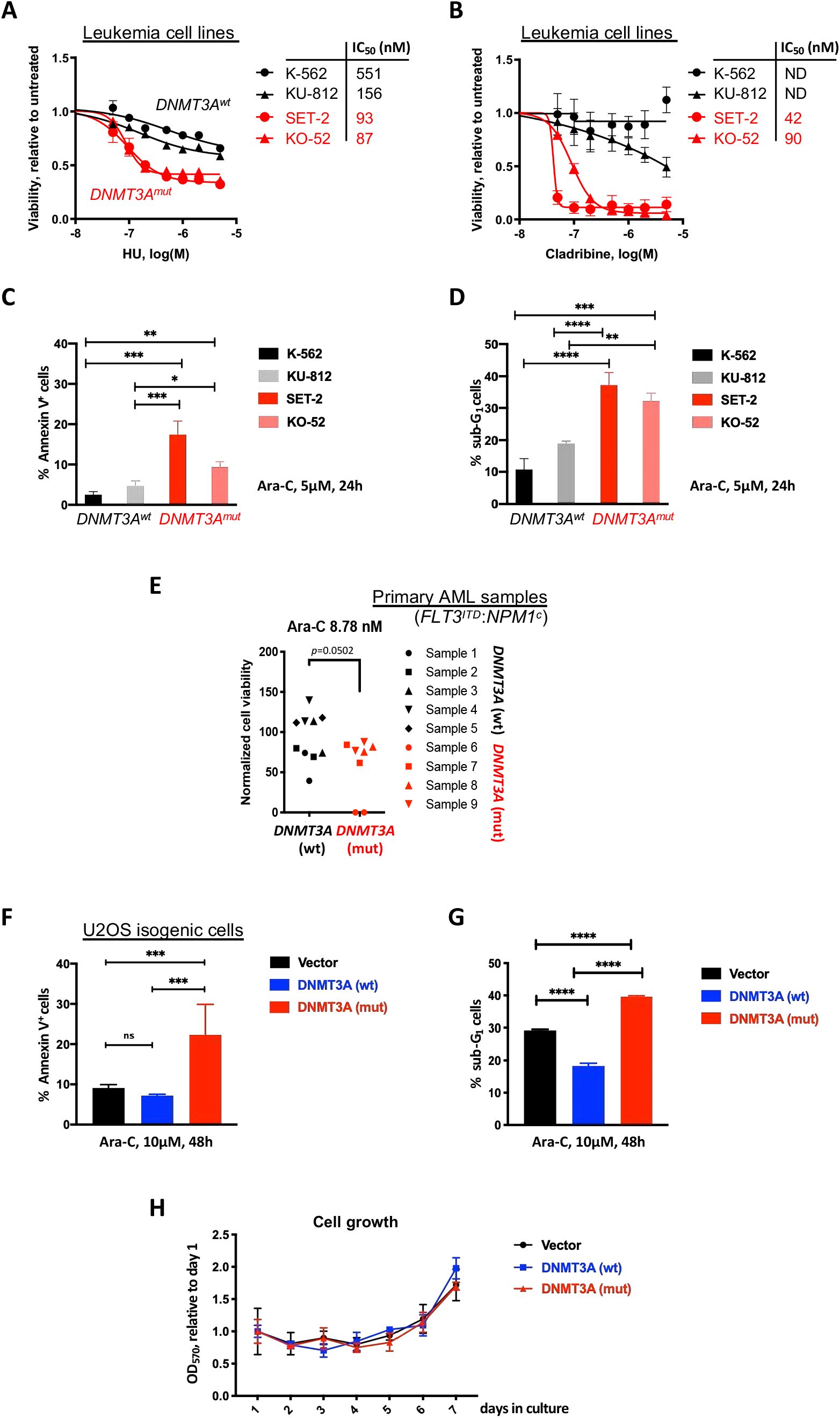
*DNMT3A* mutations confer sensitivity to replication stalling drugs. (A, B) Leukemia cells with *DNMT3A* mutations are more sensitive to replication stalling agents. Drug dose response in leukemia cells lines with wild-type *DNMT3A* (K-562 and KU-812, black) or with a *DNMT3A* R882 mutation (SET2 and KO-52, red), exposed to increasing concentrations of hydroxyurea (HU, A) and cladribine (B) for 48 hours. Cell viability was measured by CellTiter GLO assay, in triplicate and normalized to vehicle-treated controls. (C, D) Apoptosis in leukemia cell lines with wild-type or mutant *DNMT3A*, treated with 5μM cytarabine for 24 hours, as detected by Annexin V staining (C) or by DNA content stained with DAPI (D) by flow cytometry. In cell lines with mutant *DNMT3A* more cells were in sub-G_1_ compared to cells with wild-type *DNMT3A* indicative of apoptosis (* *p*<0.05, ** *p*<0.01, *** *p*<0.001, **** *p*<0.0001, two-way ANOVA with Tukey’s *post-hoc* test from three replicates). (E) *Ex vivo* cytarabine sensitivity in 5 *DNMT3A* wild-type and 4 *DNMT3A* R882 primary AML patient samples. All AMLs were *FLT3*^*ITD*^:*NPM1*^*c*^ (*p*=0.0502, two-tailed Student’s *t*-test). (F, G) Apoptosis in U2OS cells lentivirally expressing wild-type (blue) or mutant (red) *DNMT3A* or empty vector control (black) and treated with cytarabine for 48 hours, as detected by Annexin V staining (F) or by DNA content stained with DAPI (fraction of cells in sub-G_1_, G) by flow cytometry (*** *p*<0.001, **** *p*<0.0001, two-way ANOVA with Tukey’s *post-hoc* test from three replicates). (H) Similar steady-state growth rates of U2OS cells ectopically expressing wildtype (blue) or mutant (red) *DNMT3A* or empty vector control, as determined by methylene blue staining over time, relative to day 1. Relative amount of retained methylene blue was assessed by absorbance OD_570_.

**Supplementary Figure S2.**
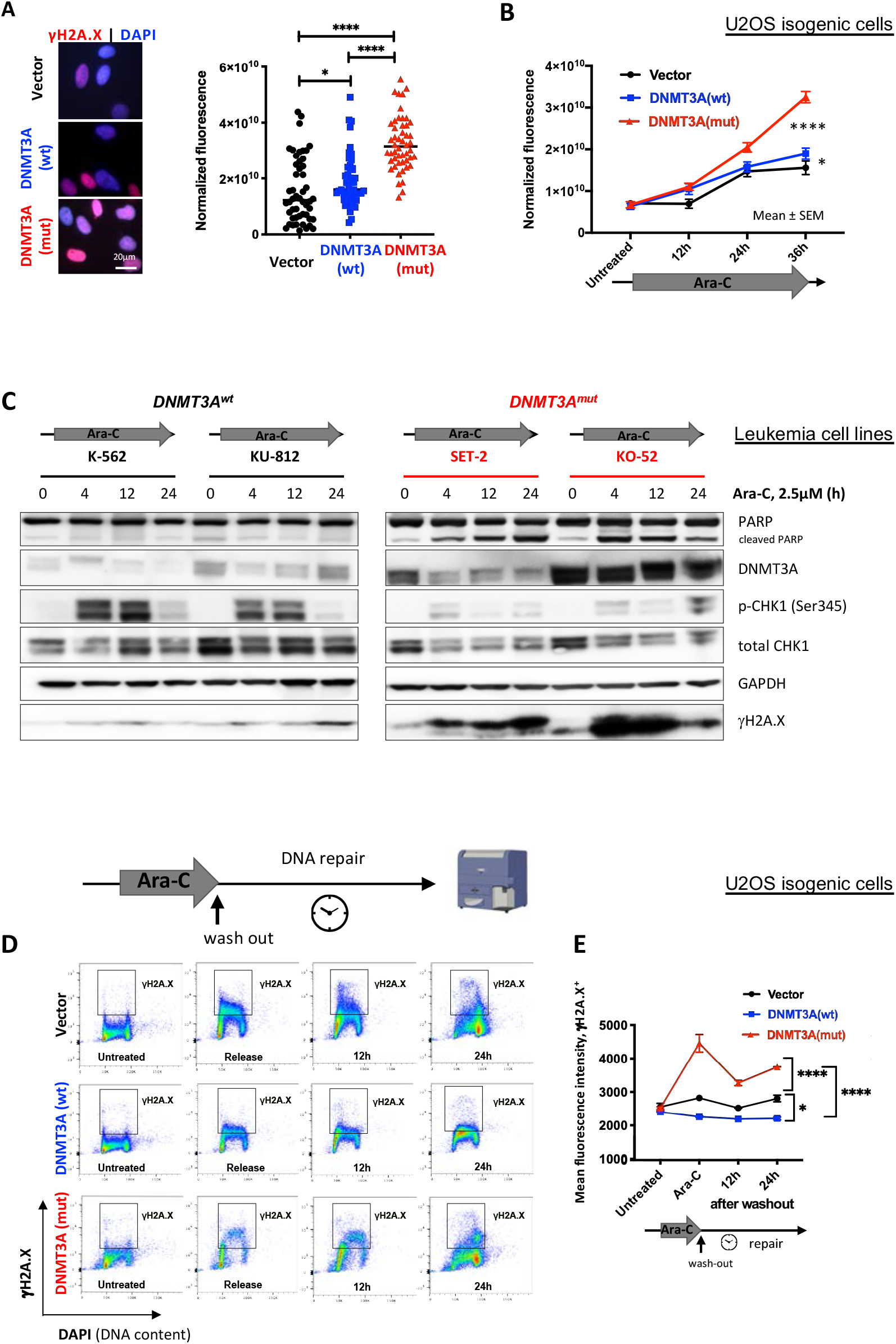
Cells expressing mutant DNMT3A accumulate DNA damage due to incomplete DNA repair after cytarabine. (A, B) Immunofluorescence staining of DNA damage marker γH2A.X 36 hours (A) or over time at indicated timepoints (B) of treatment with 10μM cytarabine, in U2OS cells lentivirally expressing wild-type (blue) and mutant (red) *DNMT3A* or empty vector control (black). For quantitative analysis, at least 50 nuclei per condition were selected at random and background-corrected fluorescence intensity was measured using ImageJ software (* *p*<0.05, *** *p*<0.001, **** *p*<0.0001, Mann-Whitney rank-sum test). (C) Immunobloting analysis of DNA damage signaling and apoptosis at indicated timepoint after cytarabine exposure (2.5μM) in cell lines with wild-type (K-562, KU-812) and R882-mutant (SET-2, KO-52) *DNMT3A*. (D, E) Persisting high levels of DNA damage in cells expressing mutant *DNMT3A* 12 and 24 hours after cytarabine wash-out. Flow cytometry scatter plots visualizing DNA damage marker γH2A.X across the cell cycle (DNA content stained by DAPI, D) and γH2A.X mean fluorescence intensity in cells surpassing DDNA damage threshold (E) in U2OS cells lentivirally overexpressing wild-type (blue) and mutant (red) *DNMT3A* or empty vector control (black), from two independent experiments performed in duplicate (* *p*<0.05, **** *p*<0.0001, Mann-Whitney rank-sum test).

**Supplementary Figure S3.**
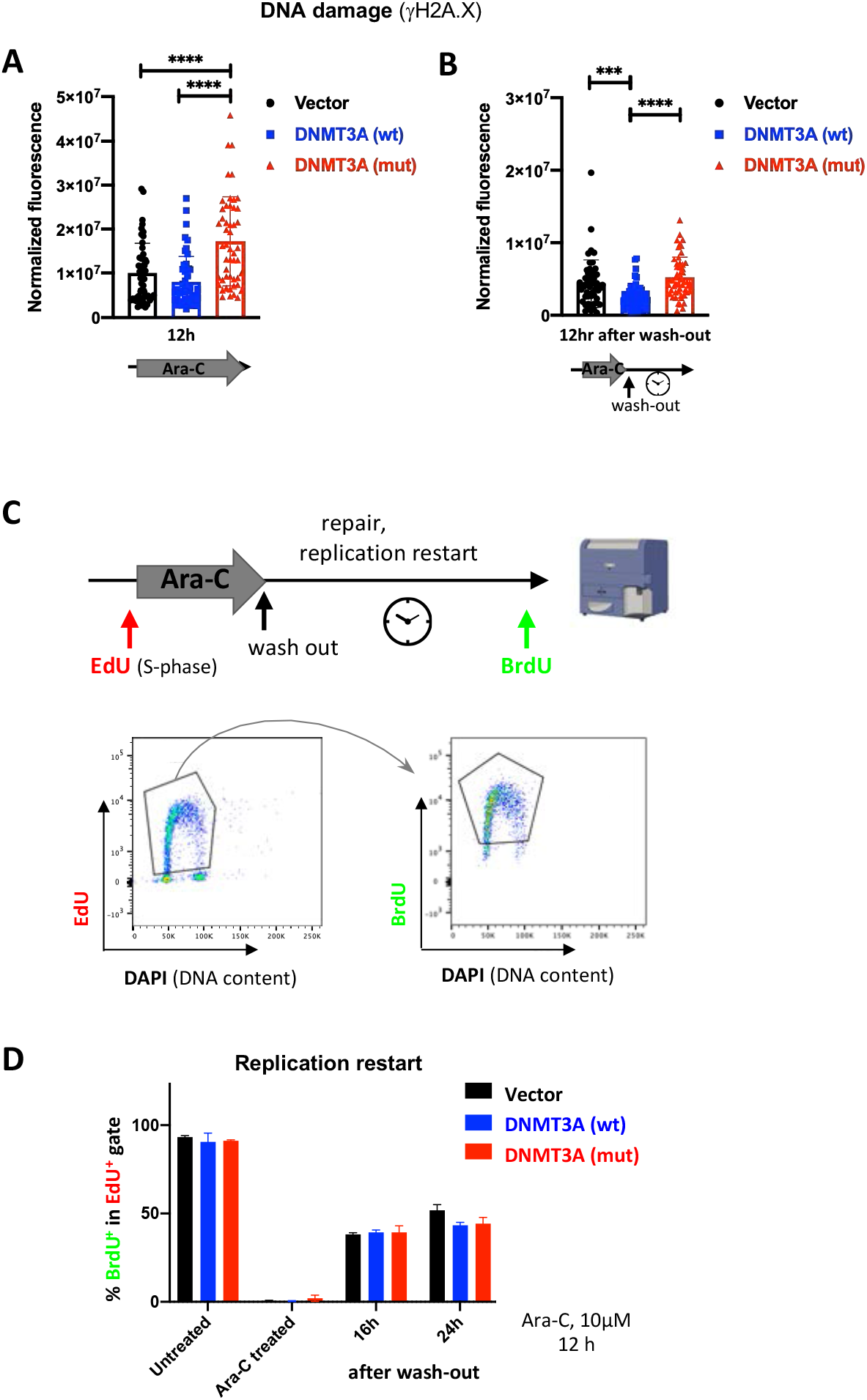
Cells expressing mutant DNMT3A accumulate DNA damage after cytarabine treatment yet are able to restart replication. (A, B) DNA damage measured by γH2A.X background-corrected fluorescence intensity after 12 hours of continuous cytarabine exposure (A) 12 hours following drug wash-out. At least 50 nuclei were quantified for each condition (*** *p*<0.001, **** *p*<0.0001, Mann-Whitney rank-sum test). (C) Experimental workflow of the replication restart analysis by dual labeling with EdU (replicating cells pre-treatment) and BrdU (at indicated time points after treatment or drug wash-out) and example flow cytometry scatter plots after exclusion of doublets and debris. Cell cycle was monitored by DNA content (DAPI). (D) Replication restart (fraction of BrdU^+^ within EdU^+^ population) in U2OS cells expressing wild-type (blue) or mutant (red) *DNMT3A* or empty vector control (black). No significant differences were detected between groups in three independent experiments each done in triplicate.

**Supplementary Figure S4.**
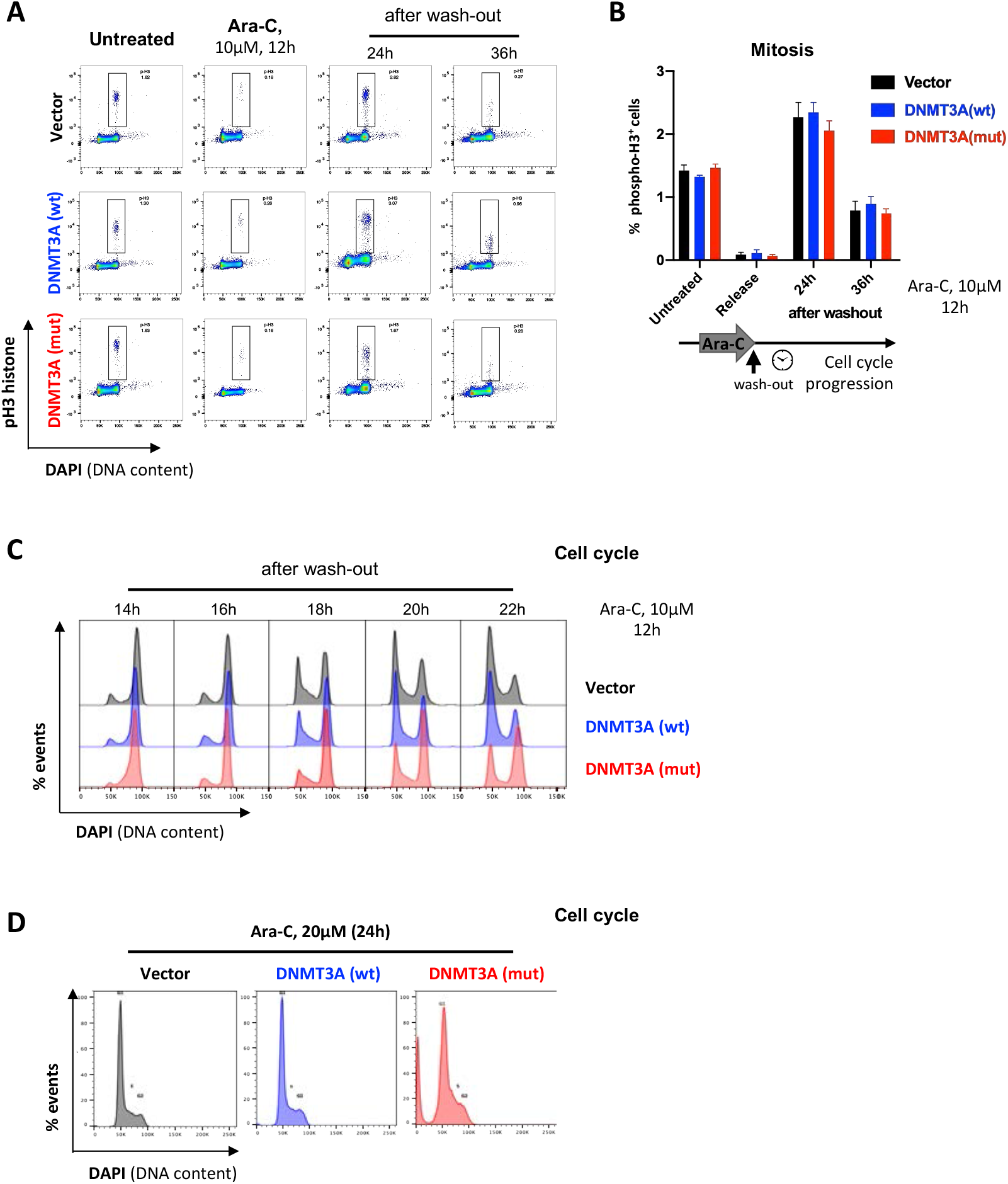
Delayed cell cycle progression in the cells expressing mutant DNMT3A after cytarabine treatment. (A, B) Progression through mitosis after cytarabine treatment and removal was determined by mitotic marker phospho-Histone H3; DAPI was used to stain for DNA content (cell cycle). Representative flow cytometry scatter plots after gating out cell aggregates and debris (A) and fraction of cells undergoing mitosis (B) after cytarabine treatment and at indicated time points after drug wash-out, in U2OS cells with wild-type (blue) and mutant (red) *DNMT3A* or empty vector control (black). (C) Cell cycle profiles (by DNA content, stained by DAPI) in U2OS cells with wild-type (blue) and mutant (red) *DNMT3A* or empty vector control (black) treated with cytarabine at indicated timepoints after wash-out. (D) Flow cytometry-based cell cycle analysis in U2OS isogenic cells after cytarabine dose intensification (20μM for 24 hours). Abnormal cell cycle profile (suppression of the G_2_ peak) in cells expressing mutant *DNMT3A* was observed.

**Supplementary Figure S5.**
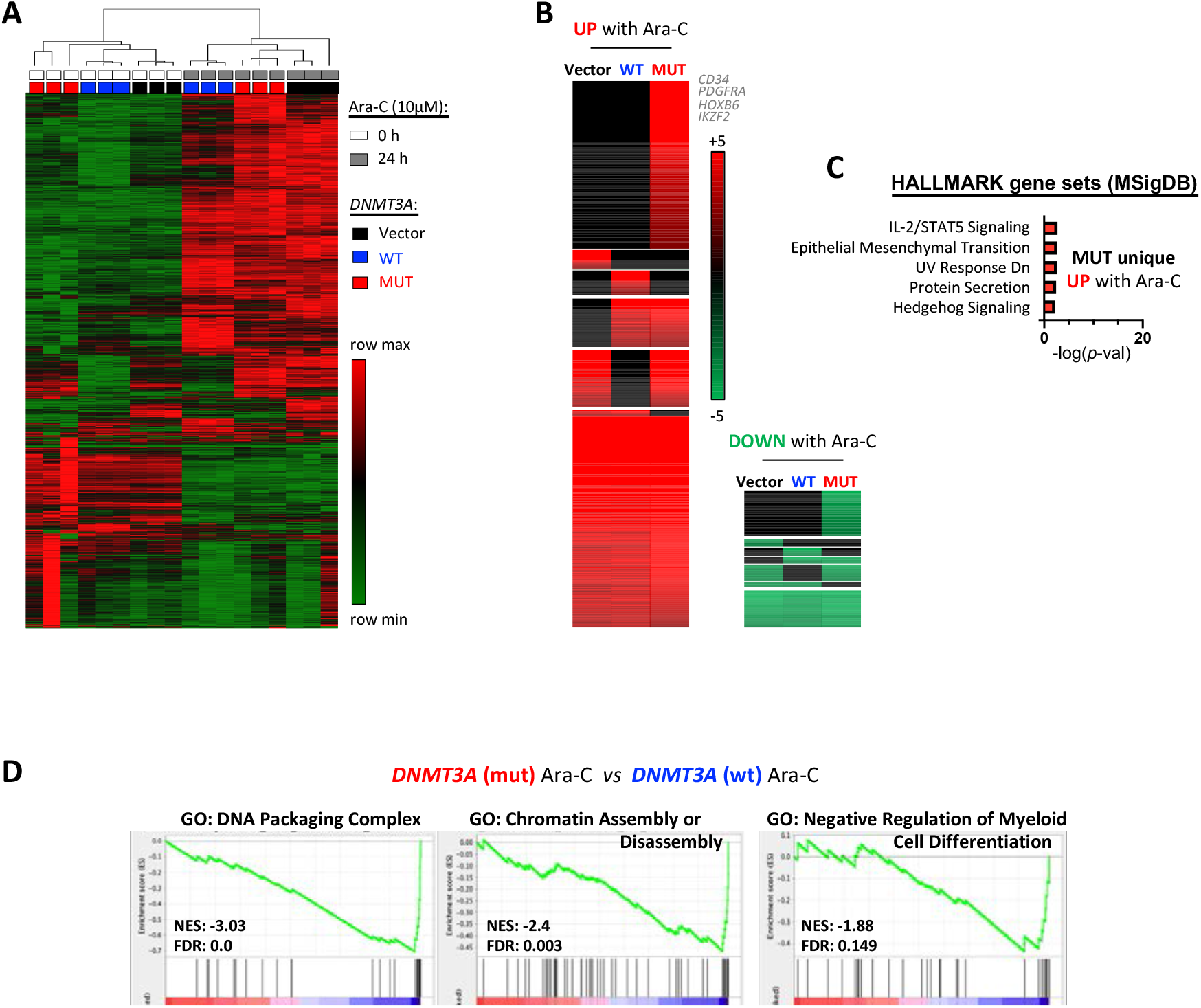
Gene expression profiling identifies pathways deregulated in cells expressing mutant DNMT3A after cytarabine treatment. (A) Unsupervised clustering on top 2000 most variable genes (by RNA-seq) demonstrates reproducibility between replicates and robust separation by treatment. U2OS cells lentivirally expressing wild-type (blue) or mutant (red) forms of *DNMT3A* or empty vector control (black) were treated with vehicle (PBS, white) or cytarabine (gray) for 24 hours, as biological triplicates. (B) Heatmap of differentially expressed genes between Ara-C-treated and untreated cells, with genes commonly or uniquely regulated in each genotype grouped together. (C) Pathways of the HALLMARK collection (MSigDB) enriched among genes uniquely upregulated in *DNMT3A*(mut) cells after cytarabine treatment (FDR<0.1). (D) Gene set enrichment analysis (GSEA) reveals negative enrichment of signatures related to chromatin organization and inhibition of myeloid differentiation in cells with *DNMT3A*(mut) compared to *DNMT3A*(wt), with cytarabine exposure.

## Supplementary tables

**Supplementary Table S1.** Primary antibodies used for immunoblotting analyses.

| Antigen | Clone | Catalog Number | Manufacturer |
| --- | --- | --- | --- |
| γH2A.X (S139) | 20E3 | #9718 | Cell Signaling Technology |
| DNMT3A | D23G1 | #3598 | Cell Signaling Technology |
| DNMT3A | C12 | sc-365769 | Santa Cruz Biotechnology |
| p-Chk1 (S345) | 133D3 | #2348 | Cell Signaling Technology |
| Total Chk1 | 2G1D5 | #2360 | Cell Signaling Technology |
| PARP | Rabbit polyclonal | #9542 | Cell Signaling Technology |
| GAPDH | 14C10 | #2118 | Cell Signaling Technology |
| p-p53 (S15) | Rabbit polyclonal | #9284 | Cell Signaling Technology |
| Total p53 | DO-1 | sc-126 | Santa Cruz Biotechnology |
| PARP1 | F-2 | Sc-8007 | Santa Cruz Biotechnology |

**Supplementary Table S2.** Primary antibodies used for intracellular flow cytometry.

| Antigen (fluorophore) | Clone | Catalog Number | Manufacturer |
| --- | --- | --- | --- |
| BrdU (FITC) | BU20A | 11-5071-42 | Thermo Fischer Scientific |
| γH2A.X<br>(AlexaFluor 647) | 2F3 | 613408 | Biolegend |
| p-Histone H3 (S28)<br>(AlexaFluor 647) | HTA28 (RUO) | 558217 | BD Biosciences |
| BrdU (AlexaFluor 488) | MoBU-1 | B35130 | Invitrogen |

**Supplementary Table S3.** Primary antibodies used for immunofluorescence.

| Antigen | Clone | Catalog Number | Manufacturer |
| --- | --- | --- | --- |
| γH2A.X | JBW301 | 05-636 | Millipore |
| PCNA | PC10 | sc-56 | Santa Cruz Biotechnology |
| p-RPA2 (S33) | Rabbit polyclonal | NB100-544 | Novus Biologicals |
| BrdU | BU-1 | MA3-071 | Invitrogen |
| BrdU | MoBU-1 | B35128 | Invitrogen |
| Lamin B1 | Rabbit polyclonal | ab16048 | Abcam |

## ^#^Abbreviations

AML: acute myeloid leukemia
ara-C: cytosine arabinoside (cytarabine)
DDR: DNA damage response
DNMT3A: DNA methyltransferase 3A
DSB: double-strand DNA break
OXPHOS: oxidative phosphorylation
PBS: phosphate buffered saline
SSB: single-strand DNA break
RT: room temperature

## Notes

### Competing Interest Statement

The authors have declared no competing interest.

